# On the overlap of transcription factor binding and regulatory targets: functional and regulatory coherence of top-bound targets is masked by weakly bound ones

**DOI:** 10.1101/2025.10.12.681120

**Authors:** Vladimir Mindel, Naama Barkai

## Abstract

A recent study that systematically mapped genomic bindings and regulatory effects of transcription factors (TFs) reported a surprisingly low overlap between TF binding and regulatory targets in *Saccharomyces cerevisiae*^1^. Our analysis suggests that this conclusion depends on the permissive definition of binding targets, which includes a large fraction of weakly bound sites.

## Results

The authors used ChEC-seq to determine TF binding locations and whole-genome nascent RNA-seq after depletion of the TFs. As we performed similar extensive binding assays and (more limited) expression assays in our lab^2-5^, but have not noticed this discrepancy, we compared our binding dataset with this new one. The datasets were highly consistent between the two groups (median genome-wide correlation between the same TFs-0.82), despite lab-specific variations in the ChEC-seq protocol. However, we noted key differences in the analysis: While the authors of the manuscript defined binding peaks (and hence binding targets) through comparison to a free MNase control^1^, a convention used in, e.g., ChIP-seq analysis, we focused on the top-bound promoters.

Tightly bound targets can be defined in several ways; we favor summing the reads mapped to each promoter and classifying promoters as bound based on their position within the resulting distribution. Using a stringent cutoff (e.g., z-score > 3.5), the number of bound promoters as predicted from the published dataset^1^ drops dramatically compared to the peaks identified by free-MNase comparison (Fig. 1A, B): instead of hundreds, only a few dozen promoters remain, and in some cases, only a handful.

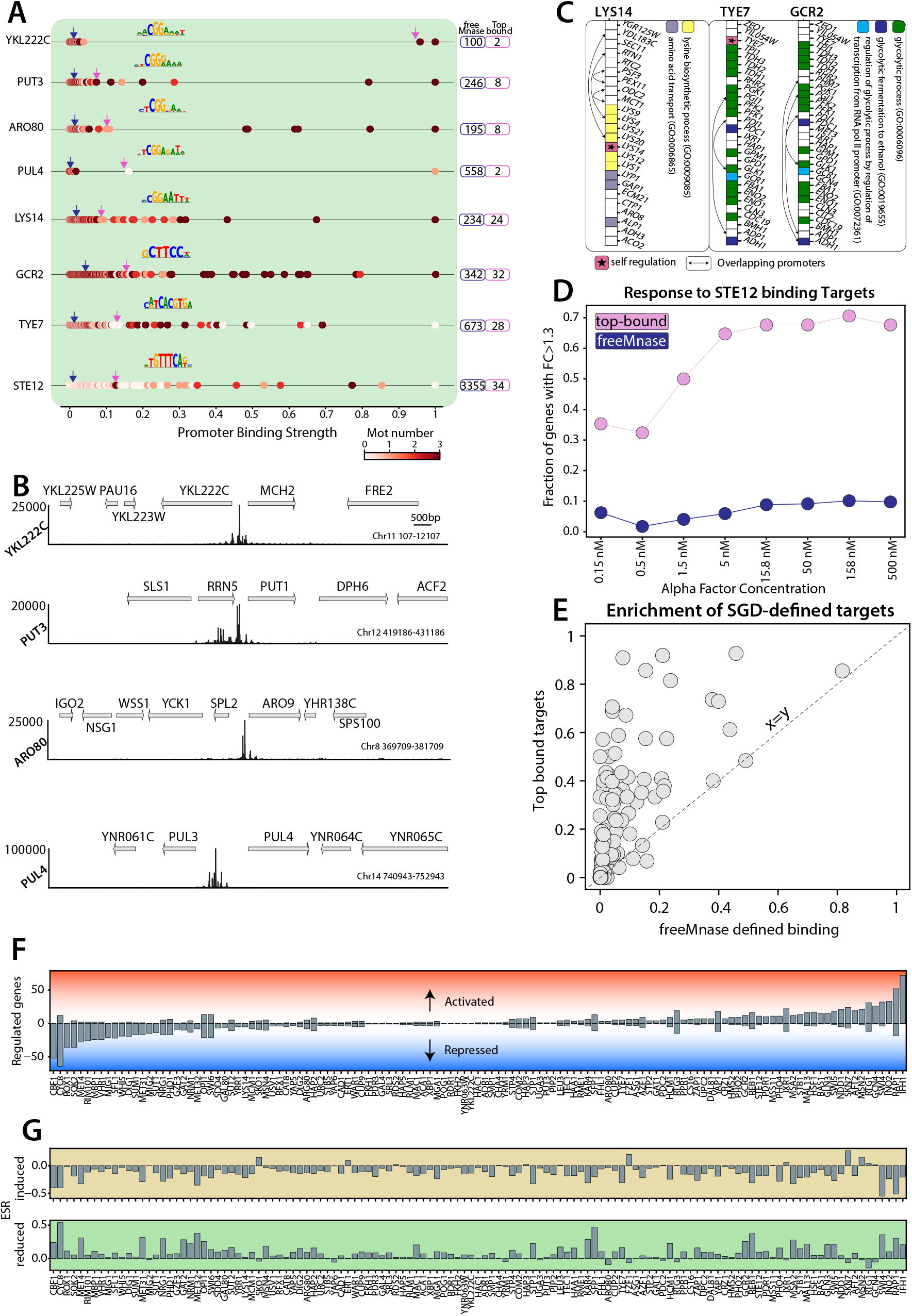
***(A) TF-binding signals vary widely across promoters***. For each promoter, the binding signal was defined as the sum of mapped reads (see Methods). Shown are relative binding signals for the indicated TFs, normalized to the strongest-bound promoter. Each dot represents a promoter, positioned along the x-axis by its normalized signal and colored by the number of motif occurrences (based on the known PWM; see Methods). Pink and blue arrows mark thresholds for defining bound targets: z-score > 3.5 (pink) or significance relative to free-MNase (blue). The numbers of targets identified by each method are shown on the right, circled in the corresponding colors. ***(B) Some TFs bind only a handful of promoters***. Shown are binding tracks for four fungal-specific zinc-cluster TFs with strong signals (z-score > 3.5) at fewer than ten promoters, contrasting 100-500 bound promoters predicted by free-MNase comparisons (see A). At this z-score>3.5 threshold, YKL222W and PUL4 each bind at a single divergent promoter (YKL22W at YKL22W /MCH2, and PUL4 at PUL3/PUT4). Note that both MCH2 and PUT4 are putative permeases. ARO80 and PUT3 display strong binding at eight promoters each, with the strongest signals at their known targets (ARO8/ARO9 for ARO80; PUT1/PUT2 for PUT3). By contrast, free-MNase comparison suggests binding at 100–500 promoters, most of which show weak binding signals. ***(C) Pathway-dedicated TFs***. Shown are strongly bound targets (z-score > 3.5) of three TFs enriched for specific metabolic pathways. Lys14 binds promoters of lysine biosynthesis genes, while GCR2 and TYE7 primarily target glycolytic genes. Bound promoters are colored by pathway: lysine biosynthesis (yellow) and glycolysis (green). Arrows indicate divergent promoters regulating two genes. Free-MNase comparison predicts 250–650 binding sites for these TFs, but most show only weak signals. ***(D) Co-regulation of top-bound promoters***. Ste12 is a known regulator of mating genes. We identified 34 genes bound at promoters with a z-score exceeding the 3.5 threshold and examined their induction by mating factor. Shown is the fraction of induced targets (see Methods) as a function of alpha-factor concentration (pink). In contrast, the free-MNase comparison predicts more than 3300 Ste12-bound promoters, which remain largely unaffected by alpha-factor. ***(E) Top-bound targets are enriched for known regulatory interactions***. For each profiled TF, we compared strongly bound targets (z-score > 3.5) to previously defined targets from the SGD database. The plot shows the fraction of SGD-defined targets among strongly bound targets (y-axis) versus among all significant targets defined by free-MNase comparison (x-axis). ***(F) Strongly bound targets display coherent responses to TF depletion***. For the set of analyzed TFs, we measured the fractions of genes activated or repressed following TF depletion (genes defined as activated/repressed if log□ fold-change exceeded ±1.3). ***(G) TF depletion elicits a stress response***. Shown are median expression values of Environmental Stress Response (ESR) genes: induced (brown) and repressed (green). These responses highlight potential indirect effects of TF depletion.

Importantly, these top-bound targets show strong functional and regulatory coherence when testing specific examples (Fig. 1B–D), and this tendency is further seen in a general analysis comparing bound promoters to know regulatory targets, as defined by public database ^6^ (Fig. 1E). For instance, Ste12 is predicted by free-MNase comparison to bind at more than 3,300 promoters, yet only 34 of these exceed the 3.5 z-score threshold. These 34 top-bound locations are highly enriched for mating-factor-responsive promoters, consistent with Ste12 being the known regulator of the yeast mating response.

More generally, top-bound targets consistently show a coherent transcriptional response to TF depletion, being either upregulated, downregulated, or unaffected in the tested condition (Fig. 1F). Note that secondary effects of TF depletion cannot be excluded in this analysis, as the environmental stress response (ESR) program is consistently activated in a large fraction of TF-depleted cells (Fig. 1G).

In conclusion, the limited overlap between TF binding and regulatory targets reported may characterize low-bound targets, while tightly bound ones are more likely to be functionally coherent and regulated by the bound TF, provided that the appropriate TF-activating conditions are met. Given these results, we call for a refined TF target definition that will yield sharper insight into transcriptional regulation in yeast and beyond.

## Methods

### Data Availability

Code and processed data are available at: https://doi.org/10.5281/zenodo.17292965

### Data processing

Raw sequencing data were downloaded from GEO (GSE272555 accession number), adaptor dimers and short reads were filtered out using cutadapt with parameters: ‘‘–O 10 –pairfilter = any –max-n 0.8 –action = mask’’. Reads were aligned to the S. cerevisiae genome R64-1-1 using Bowtie 2 with the options ‘‘–end-to-end –trim-to 40 – verysensitive’’. The genome coverage of fully aligned read pairs was calculated with genomecov from BEDTools using the parameters ‘‘-d -5 –fs 1′’. The reads were normalized to a total of 10^6, and the repetitive regions of the ribosomal rRNA genes and CUP-1/2 were excluded prior to normalization For the mean signal, the average of the normalized signal from the provided biological repeats was calculated

### Promoter boundaries definition

Transcription start sites (TSSs) were defined by comparing publicly available TSS datasets (Park et al., 2014; Pelechano et al., 2013; Policastro et al., 2020). Promoter regions (n = 5424) were then defined from the start codon until at least 700 bp upstream of the TSS (start codon if no TSS was available) or the closest verified ORF. The binding signal (bs) on each promoter was then defined as the sum of the normalized ChEC-seq signal along that promoter.

### Target promoter definition

For each TF, the sum of signal on promoters was normalized using a z-score transformation, and top-bound promoters were defined as those passing the threshold of 3.5 (z-score units). To obtain the number of target promoters defined by the comparison to the freeMnase data from the supplemental material of ^1^ was used.

### Motif analysis (Fig. 1A)

To define the number of motifs of TFs residing inside the promoter PWMs of the shown TFs, the PWMs of the TFs were downloaded from the cis-bp motif database. All promoter sequences (n=5424) were screened for the number of motif locations using the BioPython package (BioPython motifs); sequence distribution from all promoters was used as a background to the search, and the threshold of 1000 (approximately the ratio of *fnr/fpr*) was used to define the presence of motifs.

### Response to alpha-factor analysis (Fig.1D)

Gene counts of response to alpha factor from S. Cerevisiae DNA microarray data were downloaded from ^6^. A fraction of STE12 target promoters (see Target promoter definition) whose gene counts passed a log2(1.3) increase in Fold Change from the control were plotted as a function of the alpha factor concentration.

### Mining of SGD for the TF-target data (Fig. 1E)

To obtain the data on previously reported TF-target gene interactions, we analyzed data present in the SGD database. The genes marked as an interactor of the target TF were extracted. Fractions of target promoters whose genes are defined as interactors were plotted for the free Mnase (x-axis) and top-bound (y-axis) target promoter definitions.

### Analysis of the TF activator/repressor state (Fig. 1F)

Processed RNA data were downloaded from the GEO (GSE272555 accession number). Top-bound promoters were defined as activated/repressed if they passed the Fold Change (from control) of log2(1.3) in the respective direction and plotted as a barplot.

### Environmental Stress Response analysis (Fig. 1G)

The identities of the stress response genes of S. Cerevisiae (induced/repressed) were obtained from the^7^. The median change of the induced stress genes (top) or reduced stress genes (bottom) was plotted as a barplot for each TF expression data.

## Notes

### Competing Interest Statement

The authors have declared no competing interest.

